# Learning to leverage salient regions in neuro-oncology using Deep Learning

**DOI:** 10.1101/2020.10.22.350421

**Authors:** A. Grigis, A. Alentorn, V. Frouin

## Abstract

Developing an automatic tumor detector for MRI medical images is a major challenge in neuro-oncology. The availability of such a tool would be a valuable assistance for the radiologists. Numerous works have tried to segment the tumor tissues, others have attempted to localize the tumor globally. In this work we focus on this second class of methods and we compare two drastically different strategies. The first one is an assumption-free anomaly detector build over a Variational Auto-Encoder (VAE), and the second one is a VGG classifier that embed Attention-Gated (AG) units to focus on the target structures at almost no additional computational cost. This comparison is first conducted on the publicly available BraTS glioma dataset for which published performance results can serve as reference, and extended as such (ie., without transfer learning) to two internal image datasets, namely Primary Central Nervous System Lymphoma (PCNSL) and Metastasis. The results demonstrate that the VAE and AG-VGG strategies can be used, up to a certain extent, to localize brain tumors.

## 1. Introduction

Developing innovative approaches to facilitate the treatment of brain cancer patients has become a societal challenge. While some tumors such as meningioma can be easily segmented, others like glioma and glioblastoma are much more difficult to delineate. These tumors are often diffused, poorly contrasted, and can appear anywhere in the brain, in almost any shape and size. In the literature, a lot of works have addressed this segmentation task, either by using classical image processing based methods, machine learning approaches or deep learning Convolutional Neural Networks (CNNs). CNNs have been used to solve the brain tumor segmentation problem consisting of separating the different tumor tissues (active tumor, edema and necrosis) from normal brain tissues: gray-matter (GM), white-matter (WM), and cerebrospinal-fluid (CSF) [**Bakas 2019**]. A different viewpoint consists in learning to leverage salient regions useful for the tumor segmentation [**Zimmerer 2019**, **Schlemper 2019**].

In this study we focused on this second class of methods. More specifically we will apply Attention-Gated (AG) VGG [**Schlemper 2019**], and Variational Auto-Encoder (VAE) [**Zimmerer 2019**] to generate saliency or anomaly maps respectively. The ultimate goal is to leverage these maps to locate roughly brain tumors that can further be characterized by radio-genomic features [**Mazurowski 2015**]. The scope of this study is to validate these two strategies on various kind of brain tumors using two unique private datasets, namely PCNSL and Metastasis. These datasets contain Primary Central Nervous System Lymphoma, and brain Metastasis (or secondary brain tumors). In a nutshell, PCNSL is a diffuse large cell non-Hodgkin’s lymphoma of the B-cell type, restricted to the central nervous system and/or eyes without any associated systemic involvement. PCNSL accounts for 3% of all primary brain tumors. PCNSL can start in the eye known as ocular lymphoma, and may take very various forms that differ significantly from glioma or glioblastoma. As for brain metastases, they occur when cancer cells spread from their original organ to the brain. Metastatic cancer cells keep the features of the primary cancer they originated from, thus having the same name as the primary cancer. For example, breast cancer that spreads to the brain is called metastatic breast cancer, not brain cancer.

These data are extremely valuable as they offer a unique opportunity to explore a wide range of tumors subgroups. The objectives of this work is two-folds. First we aim at confirming the interest of the AG-VGG and VAE for the tumor localization on the publicly available BraTS dataset [**Menze 2015**]. More specifically, the VAE was trained following the guidelines proposed in [**Zimmerer 2019**], namely to learn the normality on HCP [**VanEssen 2012**] in order to track abnormalities on BraTS. This part of the work may be considered as a replication. As for the AG-VGG, it was initially proposed to be applied on ultrasound images [**Schlemper 2019**]. We extended this work to MRI data by training and testing the model on BraTS. Finally, we applied the two trained networks on completely new data with unfamiliar tumor types (PCNSL and Metastasis). It must be noted that no transfer learning was performed prior to the application of the trained networks. We demonstrate the interest of the AG-VGG and VAE models on our datasets.

## 2. Material and Method

### Material

This study uses the HCP [**VanEssen 2012**], BraTS 2019 [**Menze 2015**], PCNSL, and Metastasis datasets.

The HCP dataset includes young healthy subjects with T1-weighted and T2-weighted images.

The BraTS 2019 dataset includes native and post-contrast (gadolinium) T1-weighted, T2-weighted, and T2-FLAIR MRI that were collected from 19 imaging sites across the world, using distinct scanners, and protocols. All records were paired with manually-defined glioma tumor masks. Annotations comprise three parts: the contrast-enhancing active tumor, the peritumoral edema, and the necrotic and non-enhancing tumor core.

The PCNSL and Metastasis datasets are unique and relatively small, with heterogeneous data either in terms of image quality or acquisition protocols. PCNSL images were collected from 10 acquisition centers and Metastasis images concern metastases of four primary tumors types (breast, lung, melanoma and esophagus). Compared to the BraTS dataset no T2-weighted image is available on Metastasis and no T2-FLAIR images is available on PCNSL. All records were also paired with manually-defined tumor masks.

### Model architectures

The VAE architecture is described in detail in [**Zimmerer 2019**] and is available on GitHub^1^. This model is a 2D variational convolutional Gaussian encoder-decoder neural network that can be represented in a U-shape. The left portion is a successive set of down-sampling layers and the right portion is a combination of up-sampling layers. In the encoding part, the features increase by two at each level. This network has the ability to approximate data distributions by optimizing the evidence lower bound (ELBO). The hyper-parameters of the model are the number of convolutional layers (default 5), the number of initial filters (default 16), the number of latent variables (default 256), the standard deviation of p(x|z) (default 0), the image size (default 64×64) and the decoder mode (default transpose-convolution).

The AG-VGG architecture is described in [**Schlemper 2019**] and is available on GitHub^2^. This model is a 2D convolutional neural network used for classification with two Attention-Gated (AG) units. By incorporating self-gating directly in the network architecture (which is end-to-end trainable) no backpropagation-based saliency map generation is required. The network is a successive set of down-sampling layers with features increase by two at each level apart from the last layer. The AG units are located after the layers N-1 and N-2, where N is the total number of convolutional layers. The hyper-parameters of the model are the number of convolutional layers (default 5), the number of initial filters (default 8), the aggregation strategy (default fit a separate fully connected layer at each scale, and make separate predictions that are weighted averaged to get final prediction).

### Data partitioning

All HCP healthy subjects were used (~1200 subjects) in a training/validation set. BraTS data were randomly partitioned with 80% assigned to a training/validation set and 20% to a test set without mixing subjects. The test set were set aside and not used during training. The Metastasis and PCNSL data were considered as a test set only.

### Training configurations

We set all hyper-parameters to default values. We focus on T2-weighted and T2-FLAIR images only as these modalities are often used in onco-imaging protocols, and because they are sensitive to tumor edema surrounding the tumors. The VAE network was trained on HCP T2-weighted data and tested on BraTS, and PCNSL (considered globally as independent test sets, see figure 1). For the optimization we used an Adam optimizer with a mini-batch of size 16, a learning rate of 1e-3, and a scheduler that reduce the learning rate when the metric has stopped improving.

**Figure 1:**
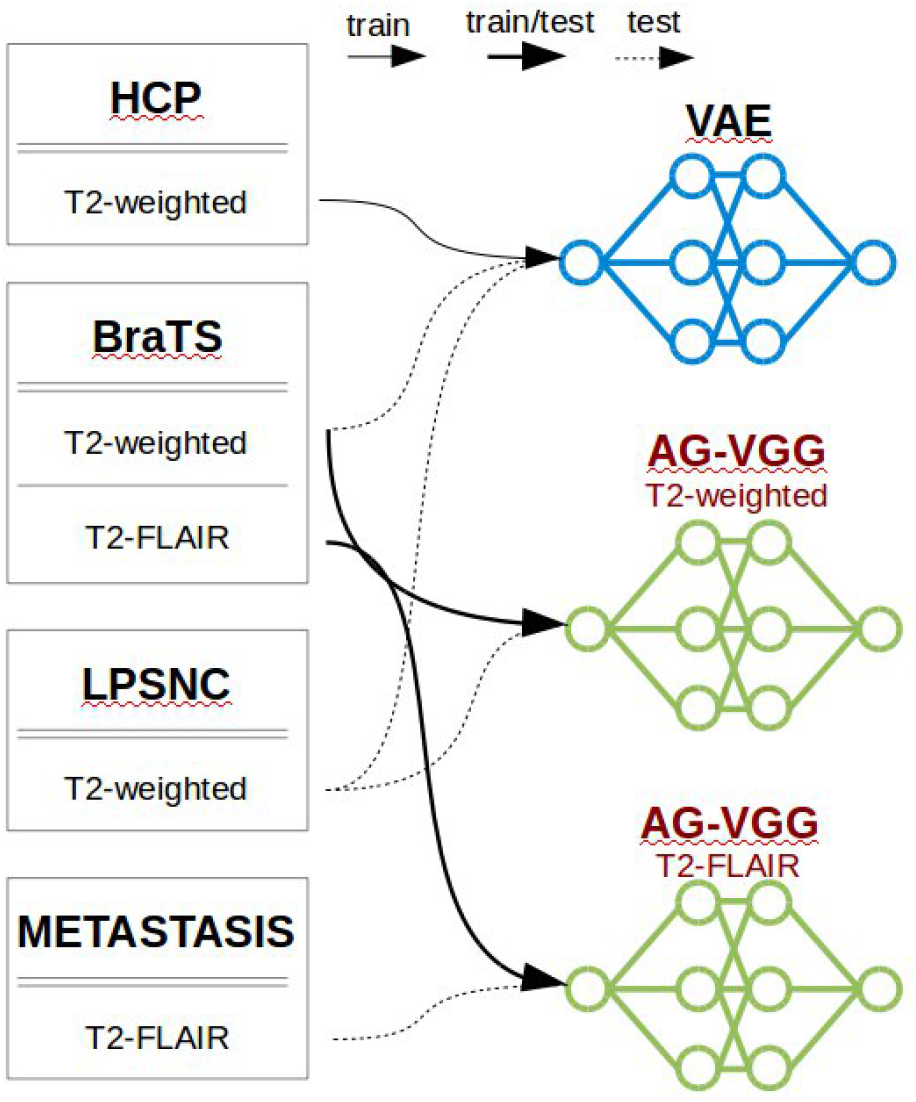
Illustration of the experimental settings.

The AG-VGG network was independently trained on the BraTS T2-weighted and T2-FLAIR data and tested on BraTS, PCNSL, and Metastasis (see figure 1). For the optimization we used an Adam optimizer with a mini-batch of size 400, a weight decay (L2 penalty) of 1e-3, a learning rate of 1e-5, and a scheduler that reduce the learning rate when the metric has stopped improving.

The assessment of the networks is achieved from the tumor segmentation Ground Truth (GT) using regular classification metrics: the precision, recall, and F1-score for each class as well as the macro average (averaging the unweighted mean per label). Tumor locations are retrieved using a simple localization heuristic that consists in i) threshold attention/anomaly maps, ii) perform connected-component analysis, iii) select the largest component, and iv) compute the smallest bounding box around the selected component.

## 3. Results

### Training: confirming the interest of tumor localization architectures on BraTS

For the VAE architecture we were able to replicate the pixel-wise performance results described in [**Zimmerer 2019**]. Briefly, we achieved an area under the receiver operator curve (ROC) of 0.72 for the pixel-wise reconstruction error (Rec-Error), 0.71 for the backpropagated ELBO (ELBO-Grad), 0.91 for its backpropagated KL-term (KL-Grad), 0.71 for the reconstruction term (Rec-Grad), and 0.87 for the Combi model. As detailed in [**Zimmerer 2019**], it seems that the reconstruction error alone is outperformed by the KL-Grad and the Combi model.

The AG-VGG architecture was originally used with ultrasound images [**Schlemper 2019**]. Here we trained the model twice with the T2-weighted and T2-FLAIR MRI data slices, and predict two labels (slice with or without tumor tissues). Table 1 summarizes the performance of the model that can detect the occurrence of tumor tissues with a precision > 0.9. In figures 2 and 3 the associated attention maps are displayed for some slices. Tumor locations are retrieved by using the localization heuristic described in the ‘Material and Method’ section. Normalized attention maps into the range [0, 1] are clear enough to fix the thresholding value at 0.5.

**Table 1:** Class-wise performance for the AG-VGG model over the BraTS, Metastasis and PCNSL datasets.

|  |  | Precision | Recall | F1-score | Support | IoU |
| --- | --- | --- | --- | --- | --- | --- |
| BraTS<br>T2w | with tumor | 0.90 | 0.82 | 0.86 | 3418 |  |
|  | without tumor | 0.87 | 0.93 | 0.91 | 4642 |  |
|  | macro avg | 0.89 | 0.88 | 0.89 | 8060 |  |
| BraTS<br>FLAIR | with tumor | 0.94 | 0.88 | 0.91 | 3418 |  |
|  | without tumor | 0.92 | 0.96 | 0.94 | 4642 |  |
|  | macro avg | 0.93 | 0.92 | 0.93 | 8060 |  |
| PCNSL<br>T2w | with tumor | 0.65 | 0.91 | 0.76 | 329 | 0.96 |
|  | without tumor | 0.89 | 0.60 | 0.72 | 399 |  |
|  | macro avg | 0.77 | 0.75 | 0.73 | 728 |  |
| META<br>STASIS<br>FLAIR | with tumor | 0.71 | 0.85 | 0.77 | 407 | 0.98 |
|  | without tumor | 0.86 | 0.72 | 0.78 | 503 |  |
|  | macro avg | 0.78 | 0.78 | 0.78 | 910 |  |

**Figure 2:**
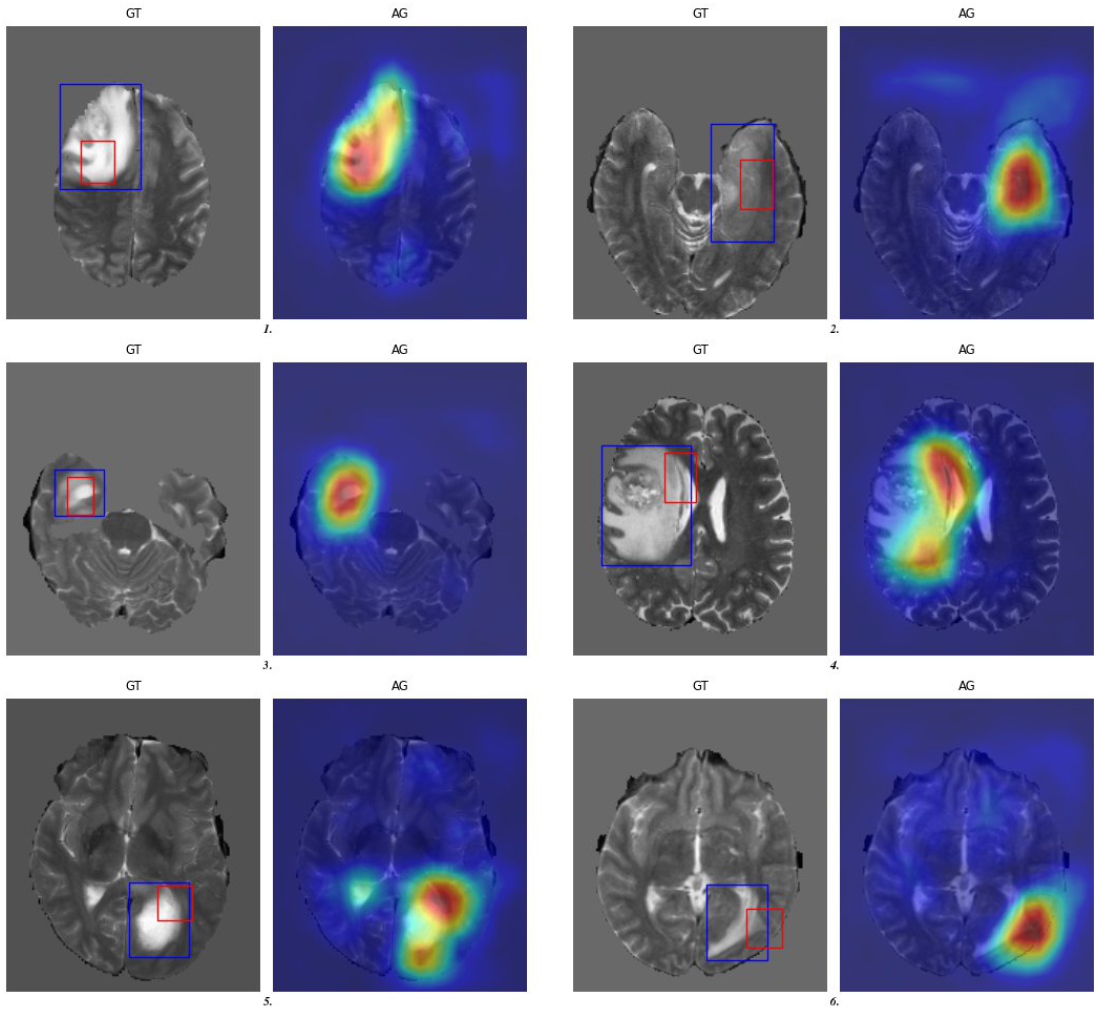
Examples of the obtained average attention maps and generated bounding boxes (red) from AG-VGG across different BraTS T2-weighted images. The ground truth annotation is shown in blue. Only true positive slices are displayed.

**Figure 3:**
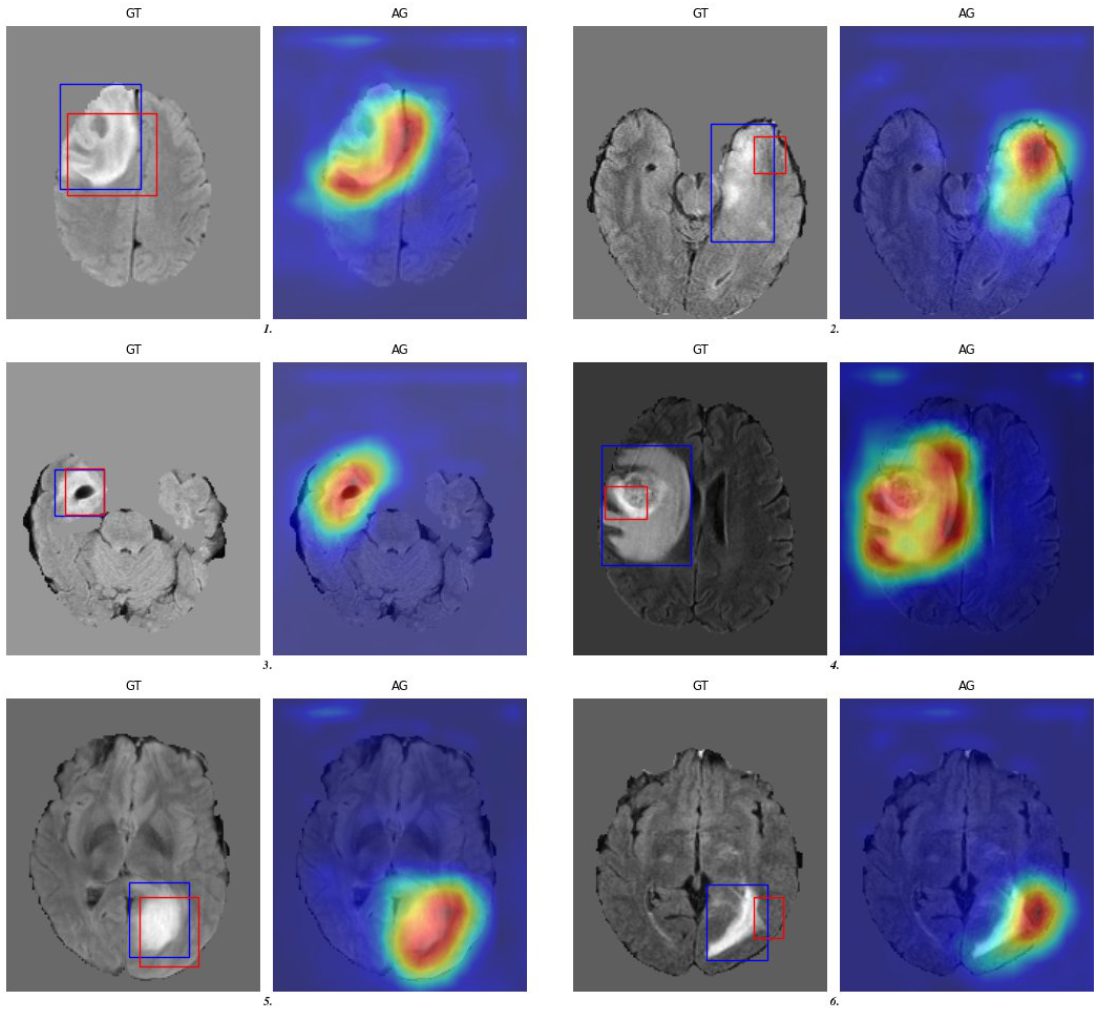
Examples of the obtained average attention maps and generated bounding boxes (red) from AG-VGG across different BraTS FLAIR-weighted images. The ground truth annotation is shown in blue. Only true positive slices are displayed.

### Application: challenging the tumor localization architectures on PCNSL and Metastasis

We applied the VAE architecture only on PCNSL as the HCP dataset does not contains T2-FLAIR images for the training (see figure 1 for the experimental plan). In figure 4 the receiver operator curves of four pixel-wise anomaly detection maps are displayed. Apart from subject d, the AUC ROC computed for the KL-Grad term is > 0.83. The AUC ROC of subject d is lower (0.75). In this particular case the network focused on a tumor removal anomaly that is prominent in the image (see figure 5). Figure 5 illustrates the detected anomaly regions that highly agrees with the GT manual tumor segmentation apart from subject d. Tumor locations are retrieved by using the localization heuristic described in the ‘Material and Method’ section. As the anomaly maps are less sharp than the attention maps, a 0.5 threshold makes no sense anymore. The thresholding becomes a complex problem. To highlight the discrimination power embedded in these anomaly maps we used the GT to control the false positive rate at 5% and select the compliant threshold. The detected region highly agrees with the GT tumor segmentation.

**Figure 4:**
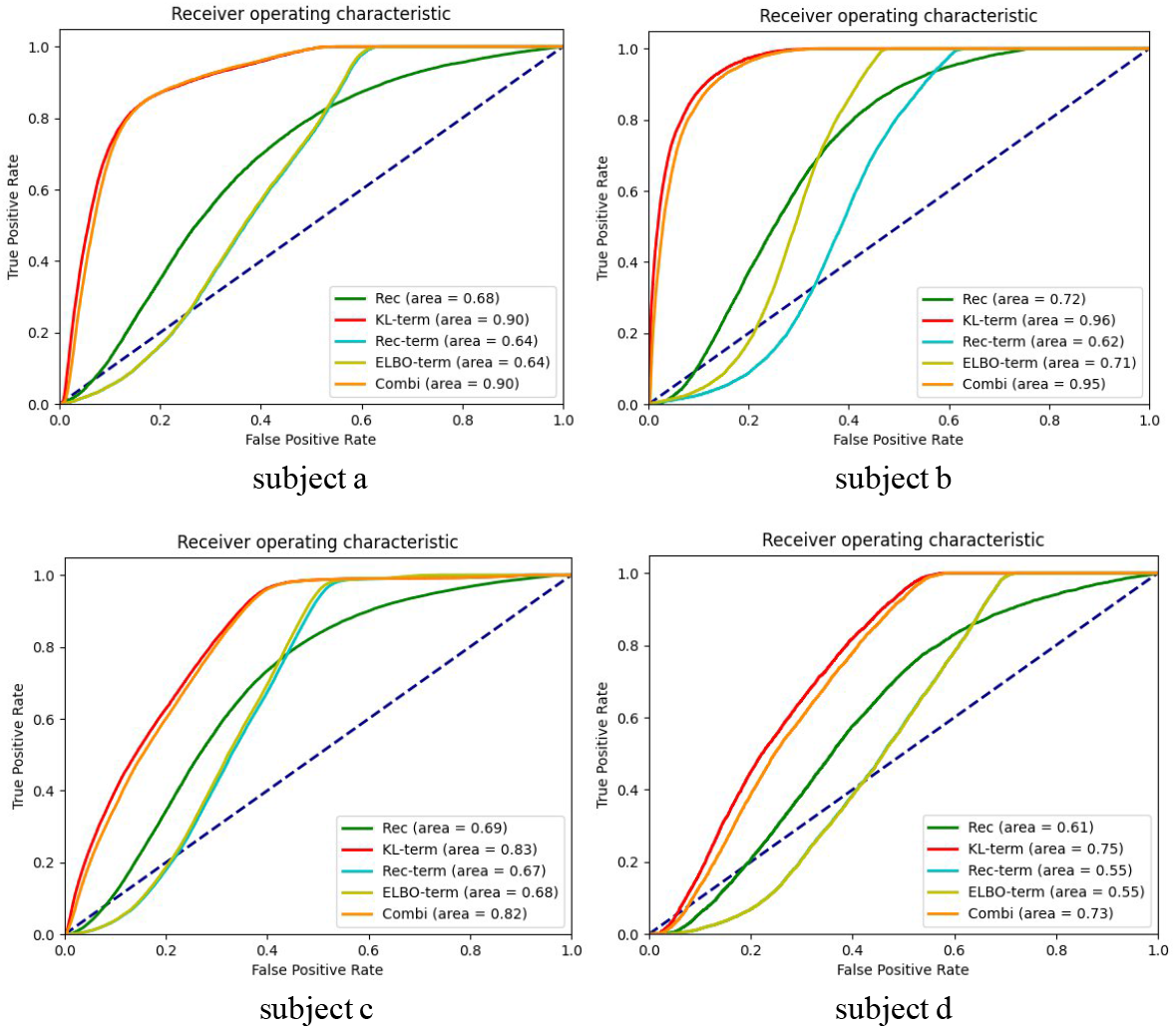
ROC curves computed from pixel-wise anomaly detection maps on four PCNSL subjects (a-d).

**Figure 5:**
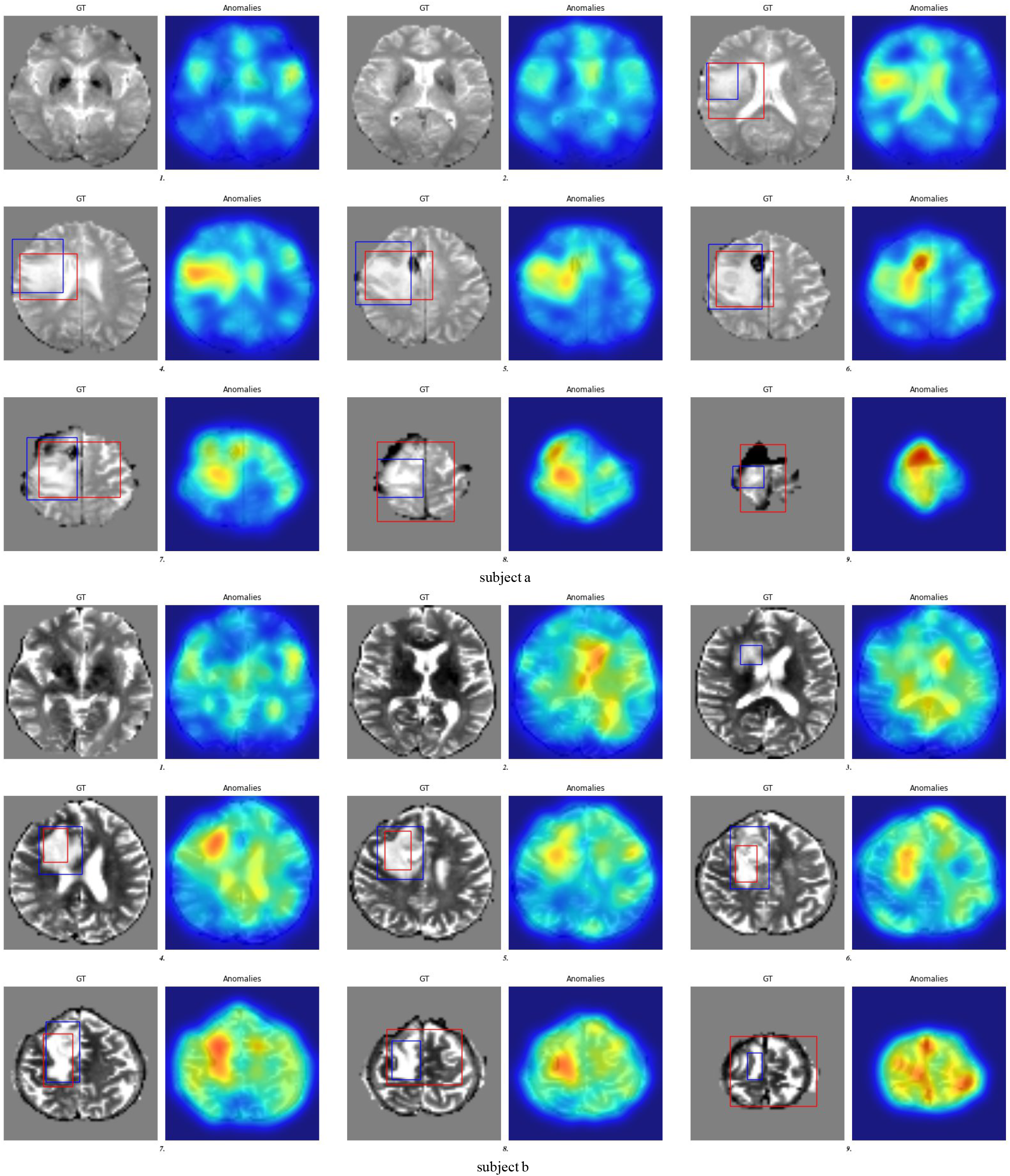
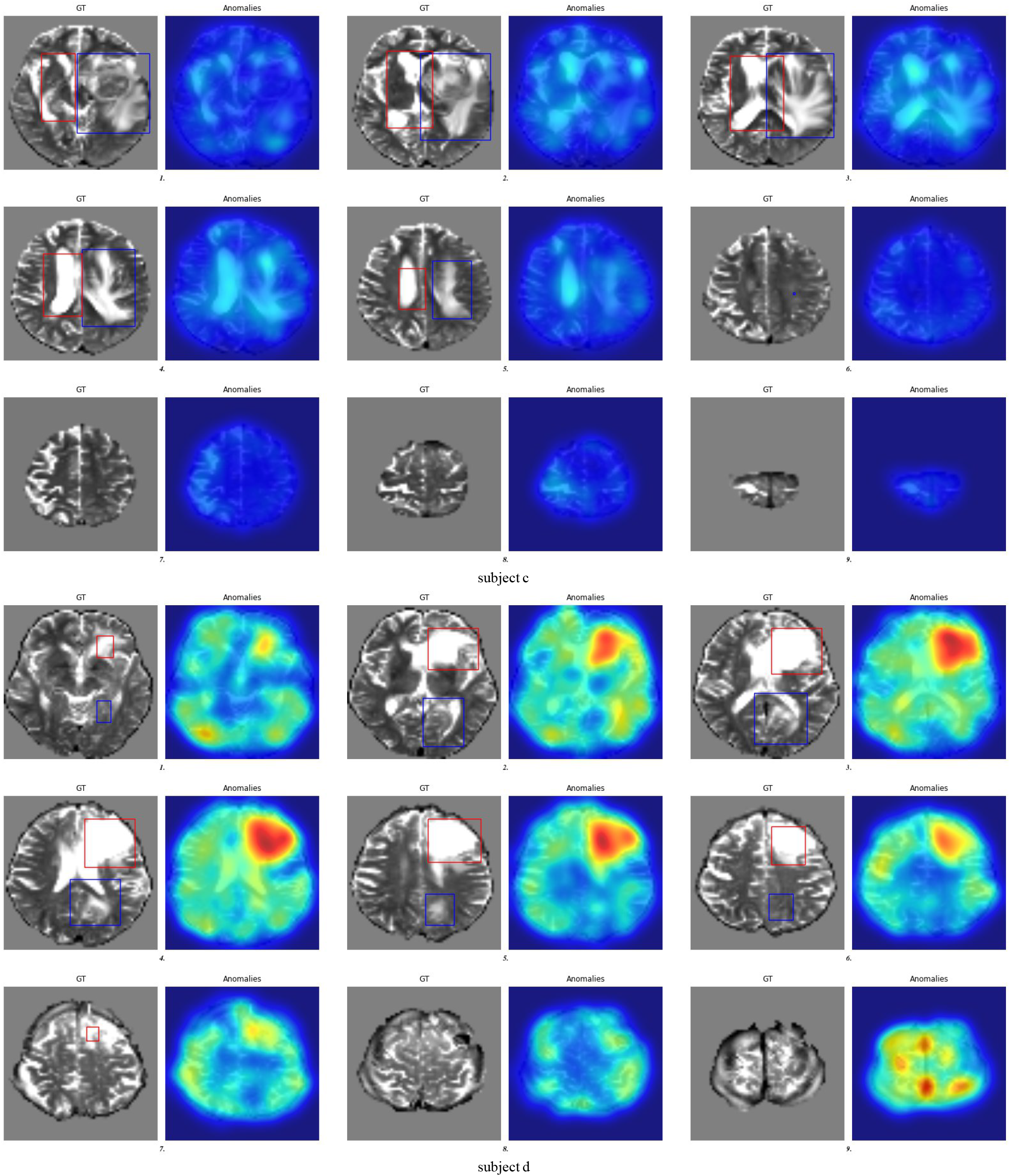
Pixel-wise anomaly detection maps on four PCNSL subjects (a-d) and generated bounding boxes (red). The ground truth annotation is shown in blue.

Then we applied the two variants AG-VGG / T2-weighted and AG-VGG / T2-FLAIR on PCNSL and Metastasis respectively. Table 1 summarizes the performance of the model that can predict the occurrence of tumor tissues with a precision of 0.65 on PCNSL and 0.71 on Metastasis. At first glance these results may seem weak. When looking at all attention maps (excluding those associated to true negative) it appears that these maps outline the discriminant tumor regions but do not completely coincide with the entire GT (see figures 6 and 7). The Intersection over Union (IoU) is a suited metric yielding 0.96 +/− 0.035 on PCNSL and 0.98 +/− 0.021 on Metastasis. We can also observe that false positive detections mainly occur because of preprocessing issues (remaining eyes, large brain mask, spikes, …) or a mix-up with the ventricles when using T2-weighted images. Note that true negatives occur even when the attention units focused on the target tumor regions.

**Figure 6:**
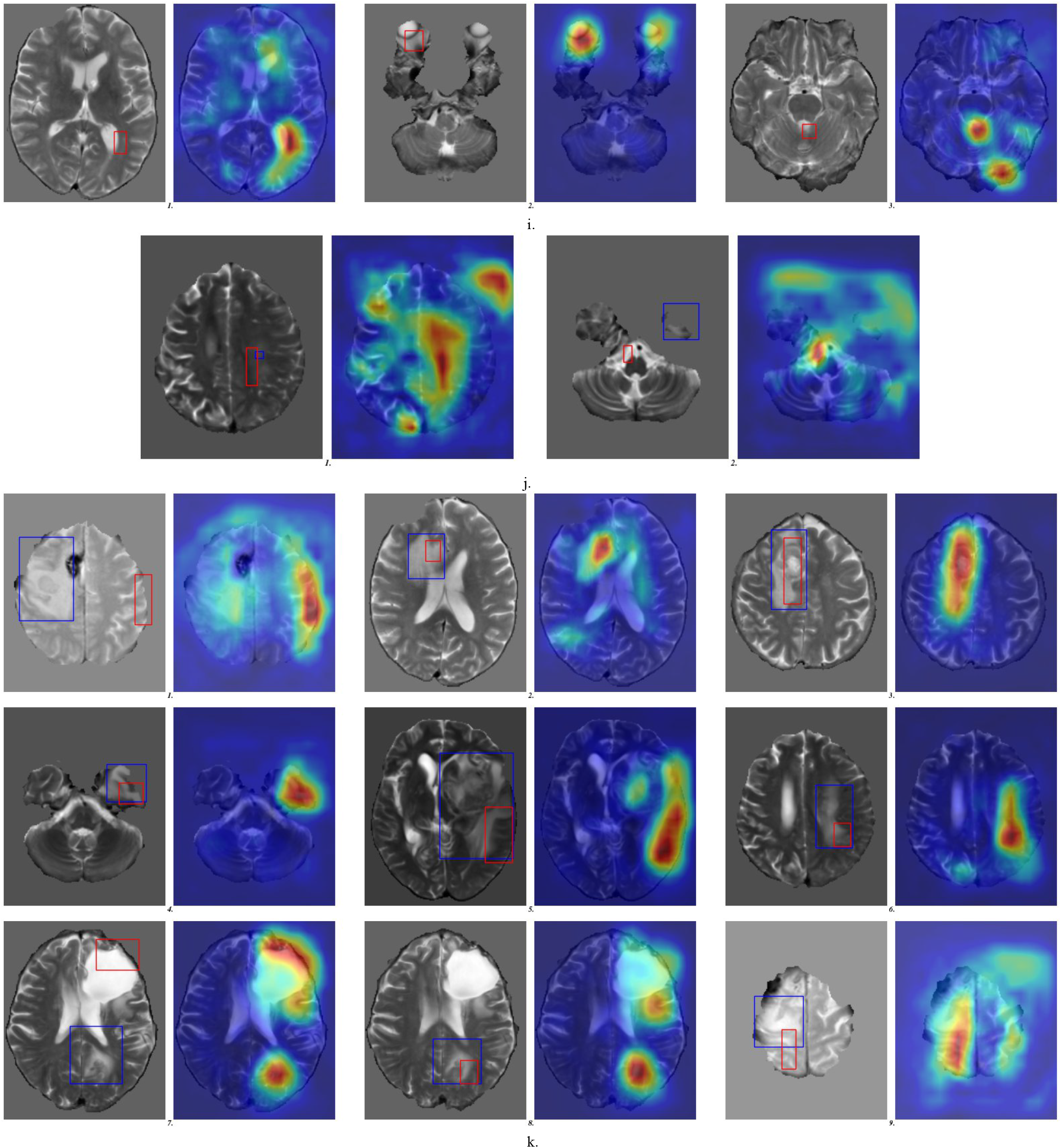
Examples of the obtained average attention maps and generated bounding boxes (red) from AG-VGG across different PCNSL T2-weighted images. The ground truth annotation is shown in blue. False positive, true negative and true positive slices are displayed (i, j, and k).

**Figure 7:**
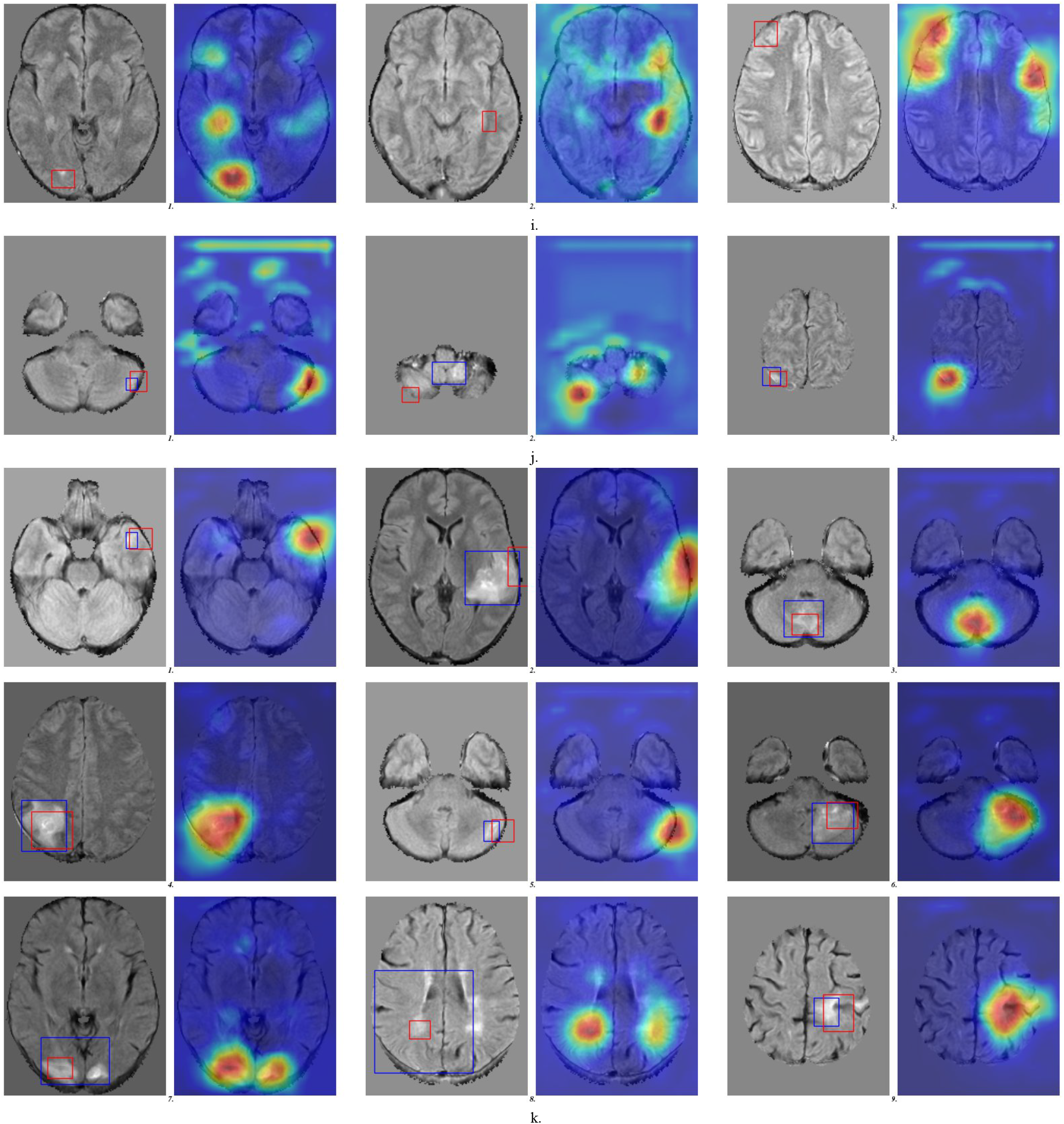
Examples of the obtained average attention maps and generated bounding boxes (red) from AG-VGG across different Metastasis T2-FLAIR images. The ground truth annotation is shown in blue. False positive, true negative and true positive slices are displayed (i, j, and k).

## 4. Discussion and Conclusion

The VAE and AG-VGG models are able to predict tumor localization with a good reproducibility and accuracy. The focused we made on 2D architectures was awarded. Indeed, the PCNSL and Metastassis data are unique, with heterogeneous data, especially with important anisotropy in the axial slice direction. Using an anomaly detection framework to locate tumors is more sensitive to false detection (ie., the tumor removal anomaly) but its genericity makes it very appealing. As for the AG-VGG, it seems to be so far the safest and most robust solution with the sharpest localization maps.

In future work we want to perform more data augmentation consistently with our data. We also believe that transfer learning may help especially for the VAE where ventricles are often targeted as anomalies. We would like to extend our subject specific study to all our data. Finally, the localization heuristic used to generate the tumor bounding box needs to be improved for instance by tracking several tumors in one slice or by using a robust thresholding. This heuristic may also take benefits of the overall 3D information.

## Acknowledgments

This work was partially funded by the PRT-K/INC a grant *LOC-model* reference 2017-1-RT-04-CEA-1.

## Footnotes

1 https://github.com/MIC-DKFZ/vae-anomaly-experiments

2 https://github.com/ozan-oktay/Attention-Gated-Networks

## Notes

### Competing Interest Statement

The authors have declared no competing interest.

### Summary of Updates

Typo in the title.

